# Transformer Enables Reference Free And Unsupervised Analysis of Spatial Transcriptomics

**DOI:** 10.1101/2022.08.11.503261

**Authors:** Chongyue Zhao, Zhongli Xu, Xinjun Wang, Kong Chen, Heng Huang, Wei Chen

## Abstract

The development of spatial transcriptomics technologies makes it possible to study tissue heterogeneity at the scale of spatial expressed microenvironment. However, most of the previous methods collapse the spatial patterns in the low spatial resolution. Existing reference based deconvolution methods integrate single-cell reference and spatial transcriptomics data to predict the proportion of cell-types, but the availability of suitable single-cell reference is often limited. In this paper, we propose a novel Transformer based model (TransfromerST) to integrate the spatial gene expression measurements and their spatial patterns in the histology image (if available) without single cell reference. TransfromerST enables the learning of the locally realistic and globally consistent constituents at nearly single cell resolution. TransfromerST firstly uses a transformer based variational autoencoder to explore the latent representation of gene expression, which is further embedded with the spatial relationship learned from adaptive graph Transformer model. The super-resolved cross-scale graph network improves the model-fit to enhanced structure-functional interactions. The public and in-house experimental results with multimodal spatial transcriptomics data demonstrate TransfromerST could highlight the tissue structures at nearly single cell resolution and detect the spatial variable genes and meta gene for each spatial domain. In summary, TransfromerST provides an effective and efficient alternative for spatial transcriptomics tissue clustering, super-resolution and gene expression prediction from histology image.

## 1 Introduction

Understanding the tissue structures at spot and subspot resolution helps to extract fine-grained information for tissue microenvironment detection. How the tissue heterogeneity shapes the structure-function interactions at enhanced resolution remains an open question in current spatial transcriptomics analysis. Modern spatial transcriptomics technologies enable to infer the large-scale structural connectivity and characterize the spatial heterogeneity patterns in disease pathology [1, 2]. The current spatial transcriptomics technologies are divided into two categories: The fluorescence in situ hybridization or sequencing based methods such as seqFISH [3, 4], seqFISH+ [5], MERFISH [6, 7], STARmap [8] and FISSEQ [9] could achieve single cell resolution. However, these technologies measure gene expression with low throughout and less sensitivity. The second category is in situ capturing based method, including spatial transcriptomics (ST) [10], SLIDE-seq [11], SLIDE-seqV2 [12], HDST [13] and 10x Visium, to measure high throughout gene expression while restraining the spatial patterns. The limitation of in situ capturing method is its low spatial resolution. The popular technologies could provide the spot measurements with 100 μm diameter in ST platform and 55 μm diameter in Visium platform. The limited resolution of current spatial transcriptomics technology requires the development of new data analysis methods to reveal the heterogeneous tissue mechanisms of tumor microenvironment, brain disorders and embryonic development [1, 14, 15].

Previous methods on spatial transcriptomics analysis could not be directly applied to link original gene expression, spatial relationship and histology image for the following reasons. 1) Most of the existing methods use dimension reduction approaches to lower the computational complexity. However, the reduced features violates the heterogeneity in gene expression in some tissues. 2) Some workflows such as Seurat [16] is developed for single cell RNA-seq analysis and corrupts the spatial relationships. 3) As far as we know, little efforts has been made to study the heterogeneity across tissue structures in both spot and enhanced resolution. Several approaches such as RCTD [17], stereoscope [18], SPOTlight [19], SpatialDWLS [20], and cell2location [21] have been presented to integrate the single cell RNA-seq with spatial transcriptomics to enhance the spatial gene resolution. However, these kinds of methods require the availability of suitable single cell reference. In many cases, the single-cell references are not available due to the limitations of budgetary, technique, and biological issues [22, 23]. Some deconvolution methods use public single-cell RNA-seq references such as Human Cell Atlas [24], BRAIN Initiative Cell Census Network (BICCN) [25], and Human BioMolecular Atlas [26] to solve the problem, but the batch effects and tissue heterogeneity in samples may result in incomplete cell types. Moreover, single-cell references and spatial transcriptomics are affected by different perturbations, which may affect the deconvolution accuracy [27].

None of the previous spatial transcriptomics analysis methods could enhance the gene expression to single cell resolution without the usage of single cell RNA-seq data. BayesSpace [28] utilizes a Bayesian prior to explore the neighborhood structure and increase the resolution to subspot level, which is coarse than single cell resolution. However, the highly computational complexity and lack of flexibility hinders its application in multimodal spatial transcriptomics data analysis. CCST [29] applies graph convolutional networks to combine the gene expression with global spatial information. SpaGCN [30] combines gene expression, spatial information and histology image through a graph convolution model. Notably, most of the existing methods such as BayesSpace, CCST and SpaGCN rely on principle component analysis (PCA) to extract the highly variable features, which is not applicable to explore the nonlinear relationships. STAGATE [31] adopts an adaptive graph attention auto-Encoder to identify spatial domains. It achieves better performance for the identification of tissue types and highly expressed gene patterns. However, the utility of STAGATE is limited to spot resolution analysis. StLearn [32] uses deep learning method for the image domain and uses linear PCA to extract the features of spatial gene expression. The lack of consideration of gene expression and spatial relationship hinders its performance in different platforms. STdeconvolve applies latent Dirichlet allocation (LDA) to deconvolve proportional representation of cell type in each multi-cellular pixel. However, STdeconvolve [33] may fail to deconvolve distinct cell type if there is no highly co-expressed genes for each cell type. STdeconvolve could not identify the location of each cell type within each multi-cellular pixel.

To address these issues, we develop a novel Transformer based framework (TransformerST) for associating the heterogeneity of local gene expression properties and revealing the dependency of structural relationship at nearly single cell resolution.TransformerST consists of three components, the conditional transformer based variational autoencoder, the adaptive graph Transformer model with multi-head attention, and the cross-scale super-resolved gene expression reconstruction model. The first component takes together, transformer and convolutional architectures to model the realistic local gene expression patterns in an effective and expressive way. The convolutional model learns the context-rich codebook with the gene expression. The long-range spatial interactions is included using the transformer architecture which models the indices distribution with a conditional constraints. The adaptive graph transformer approach identifies the tissue types with the integration of spatial gene expression, spatial relationship and histology image. It also utilizes an adaptive parameter learning step to better explore the relationship between spatial gene features and graph neighboring dependence. The super-resolved resolution is enhanced through the cross-scale internal graph neural network, which recovers more detailed tissue structures at the nearly single cell resolution. The proposed method has the following advantages,

- The proposed method provides insights into the spatial transcriptomics structural-functional dynamical relationship at nearly single cell resolution. Although the integration of single-cell RNA-seq data is widely used in deconvolution research [34–36], it may introduce bias when single-cell measurements is not available for real world applications. The proposed method does not require the single-cell RNA-seq data to infer tissue microenvironment at spot and nearly single cell resolution.
- The proposed method makes it possible to incorporate the heterogeneous spatial gene expression with histology image using multimodal data. While most of the existing methods utilize the linear PCA for feature extraction, the proposed method learns and reconstructs the original expressive gene pattern with a large number of highly variable genes (HVGs). The proposed method provides a novel pipeline for tissue type identification, spatial-resolved gene reconstruction and gene expression prediction from histology image (if available). It can be easily transferred to different spatial transcriptomics platforms, such as ST or 10x Visium.
- The proposed method is evaluated with the meta-analysis to explore the relevance of different tissue types and characterize the complex cell-cell interactions into nearly single cell resolution. The proposed method is the first time to reconstruct the gene expression at nearly single cell resolution without the usage of single cell RNA seq reference. The experimental results with different spatial transcriptomics data demonstrate the efficiency and effectiveness of proposed method to achieve better representation than state-of-the-art methods.

## 2 Results

### Overview of the proposed method and evaluations

The workflow of the proposed method is shown in Fig. 1. The key problem in spatial transcriptomics analysis is to detect the spatial patterns of gene expression. In order to learn and utilize the spatial information in spatial transcriptomics, we adopt the Graph transformer, which has great potential to link spatial information to spatial graphs. The proposed method first learns the nonlinear mapping through variational encoder component (Fig. 1a). The variational encoder helpes to explore the gene expression pattern within each spot. Simultaneously, the adaptive graph transformer is utilized to aggregate the gene expression using the corresponding neighbors relationship and histology image (Fig. 1b). The gene representation and spatial embedding are concatenated to reconstruct the original gene expression through the decoder component. The iterative unsupervised deep clustering model is introduced to detect the heterogeneous tissue types at the original spot resolution. The adaptive graph transformer enables to associate the spatial patterns with gene expression at spot resolution. To further enhance the spatial gene expression resolution, the cross-scale internal graph networks takes the concatenated embedding and histology image (if available) as the inputs to synthesize the gene expression at the nearly single cell resolution (Fig. 1c). Finally, the conditional transformer architecture (Fig. 1a) is employed to enhance the compactness of learned latent representation with the conditional constraints.The conditional transformer further explores the spatial gene expression patterns to reconstruct the corresponding codebooks.

**Fig. 1.**
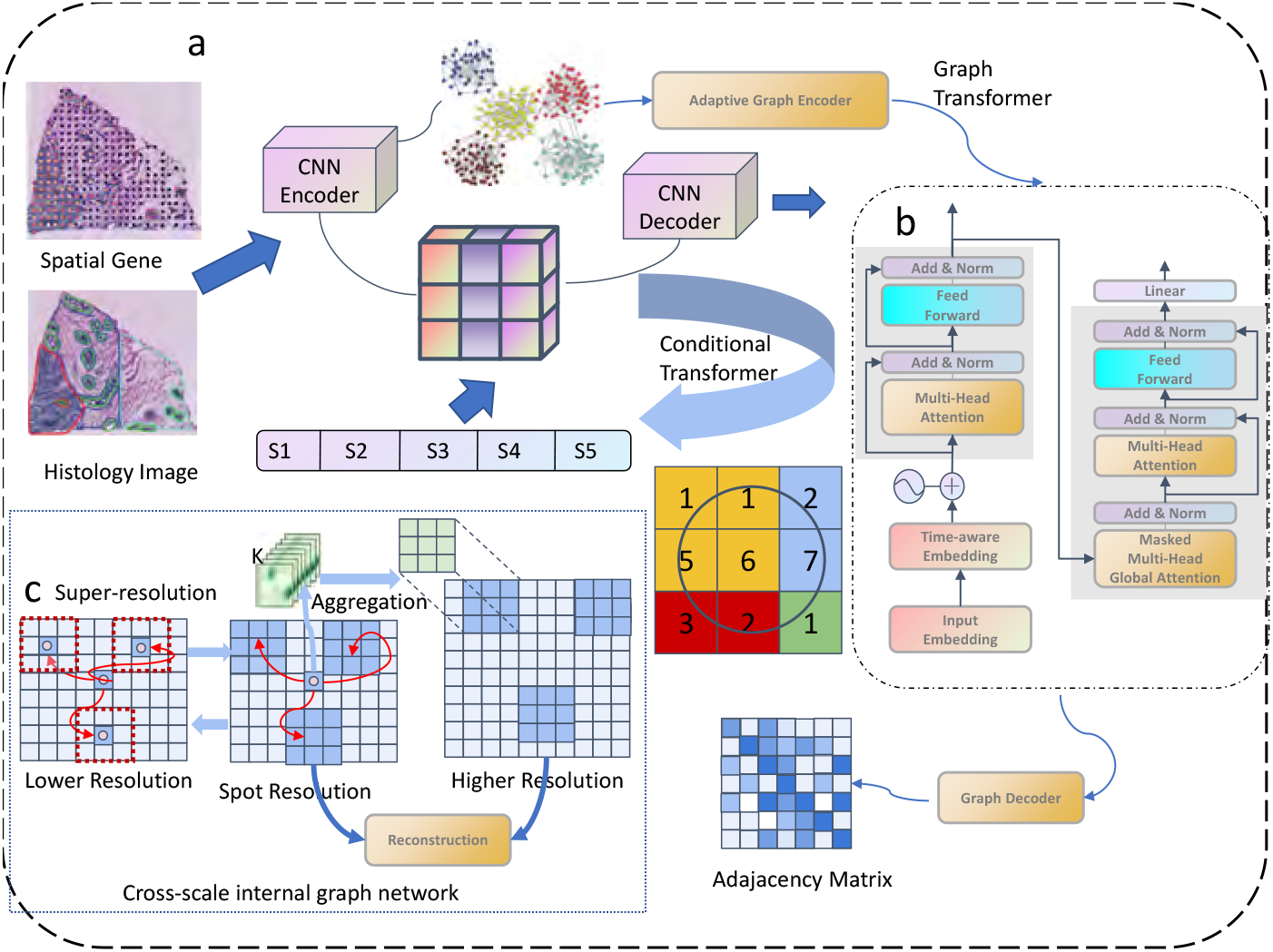
Schematic illustration of TransformerST. a, Conditional Transformer based variational autoencoder to integrate the spatial gene expression, spatial location and histology image. The variational encoder helped to explore the gene expression pattern within each spot. The conditional transformer further explores the spatial gene expression patterns to reconstruct the corresponding codebooks. b, Adaptive Graph transformer model to exploit the spatial neighboring dependence. The adaptive graph transformer model enables to associate the spatial gene expression patterns at the original resolution. c, Cross-scale internal graph network for super-resolved gene expression reconstruction. The cross-scale internal graph networks takes the concatenated embedding and histology image as the inputs to synthesize the gene expression at the single cell resolution.

To showcase the strength of the proposed method, we evaluated its performance with several publicly available datasets. In tissue identification experiments at original resolution, we showed the spot resolution clustering results with human dorsolateral prefrontal cortex data (DLPFC). We further verified TransformerST using our in-house mouse lung data with 10x Visium platform (Fig. 2 and Fig. 3). TransformerST outperformes several state-of-the-art approaches such as stLearn [32], Mclust, Kmeans, Louvain, Giotto, BayesSpace [28], CCST [29], STAGATE [31] and SpaGCN [30]. To evaluate the super-resolution performance of TransformerST, we used three data from different spatial transcriptomics platforms. Specifically, we used the melanoma data from ST platform to evaluate the super-resolution performance at subspot resolution when the histology image is missing (Fig. 4). We used human epidermal growth factor receptor(HER) 2 amplified (HER+) invasive ductal carcinoma (IDC) acquired using 10x Visium platform to show the enhanced resolution performance at nearly single-cell resolution (Fig. 4). The invasive ductal carcinoma were manually annotated by a pathologist to exclude the overexposed regions. Moreover, we also used the 36 tissue sections from the HER2+ breast cancer data to demonstrate the performance of gene expression prediction and super-resolution of TransformerST (Fig. 5). We conducted two types of experiments: the leave-one-out evaluation (36 fold) and single section evaluation. Specifically, for leave-one-out evaluation, we used 32 sections to train the clustering and super-resolution model and used the remaining section for evaluation (TransformerST). We also showed the clustering results of TransformerST using single tissue section (*TransformerST*\*). We further evaluated the super-resolution performance at nearly single cell resolution (Super-resolution). Next we investigated the spatial variable genes (SVGs) and meta gene detection accuracy using DLPFC and IDC samples. Notably, the proposed method could lower the computational complexity and reconstruct the enhanced gene expression at nearly single cell resolution more efficiently. The spatial variable genes (SVGs) and meta genes detected by the proposed methods show better biologically interpretability (Fig. 6). It should be noted that all the baseline methods were applied with the default parameters.

**Fig. 2.**
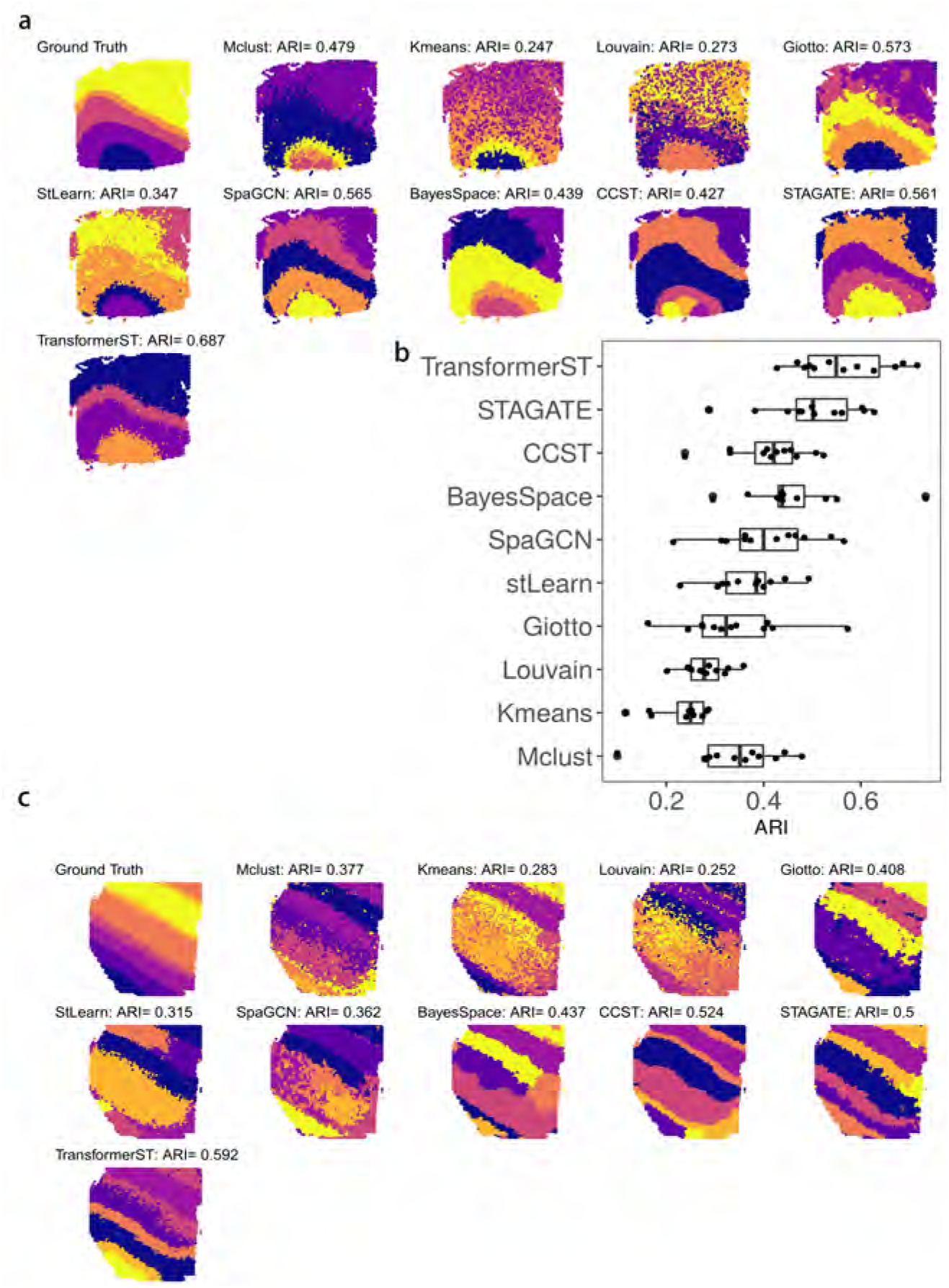
Tissue identification in human dorsolateral prefrontal cortex Visium data at spot resolution. The ARI is used to evaluate the similarity between cluster labels acquired by each method against manual annotations. a, Tissue types assignments by different spatial clustering methods for sample 151672. b, Summary of all 12 samples clustering accuracy. c, Tissue types assignments by different spatial clustering methods for sample 151508.

**Fig. 3.**
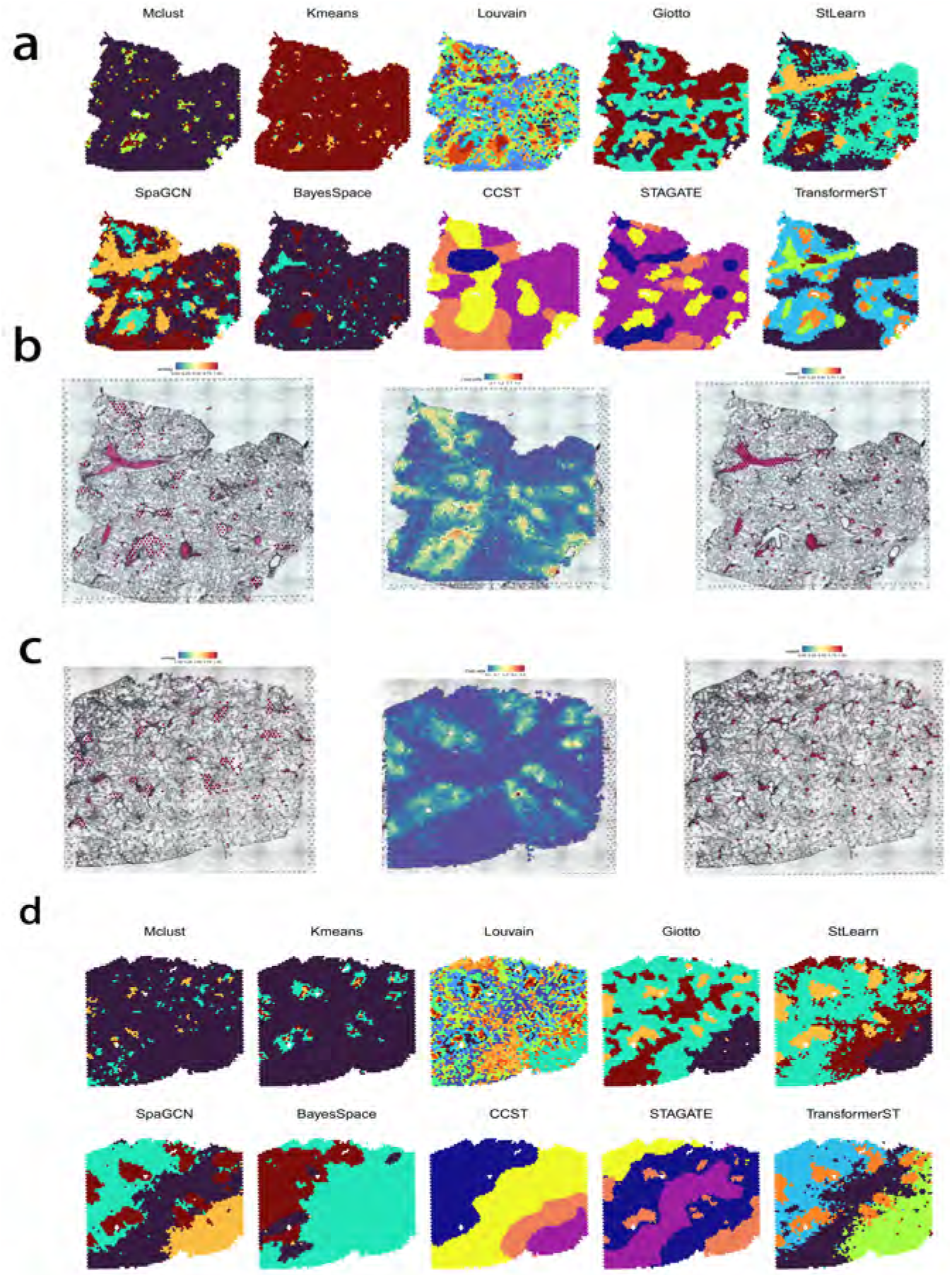
Tissue identification in mouse lung Visium data at spot resolution. a, Tissue types assignments by different spatial clustering methods for the first sample. b, Manual annotations of airways (left) and blood vessels (right) of the first slice. Pathologist identified regions of significant regions according to the histology image. Airways were defined in line with the proportion of club cells (middle) within each slice. c, Manual annotations of airways (left) and blood vessels (right) of the second slice. d, Tissue types assignments by different spatial clustering methods for the second sample.

**Fig. 4.**
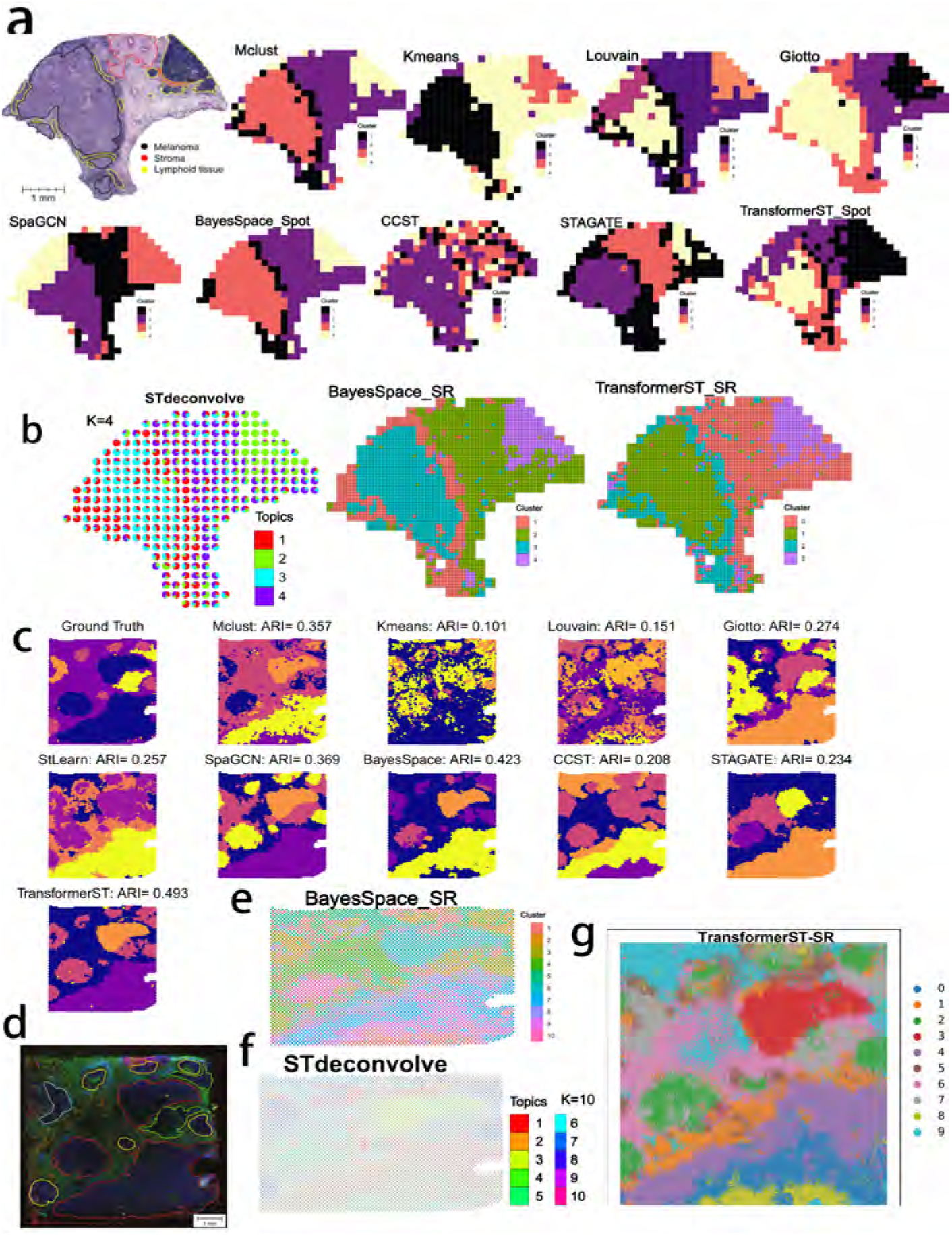
Tissue identification with super-resolved gene expression. a, Tissue type assignments by different spatial clustering methods for melanoma sample. b, Enhanced subspot tissue identification of melanoma sample with BayesSpace, STdeconvolve and TransformerST. c, Tissue type assignments by different spatial clustering methods for IDC sample. d, Immunofluorescent imaging of tissue and manual annotations. Different tissue types are shown in different colors (DAPI intensity in blue, anti-CD3 intensity in green, the Visium fiducial frame in red). Pathologist annotated different regions in different colors (IC outlined in red, carcinoma in yellow, benign hyperplasia in green, unclassified tumor in grey). e, Enhanced super-resolved tissue identification of IDC sample with BayesSpace at subspot resolution. f, Cell type proportion of IDC sample with STdeconvolve. g, Enhanced super-resolved tissue identification of IDC sample with TransformerST at nearly single cell resolution.

**Fig. 5.**
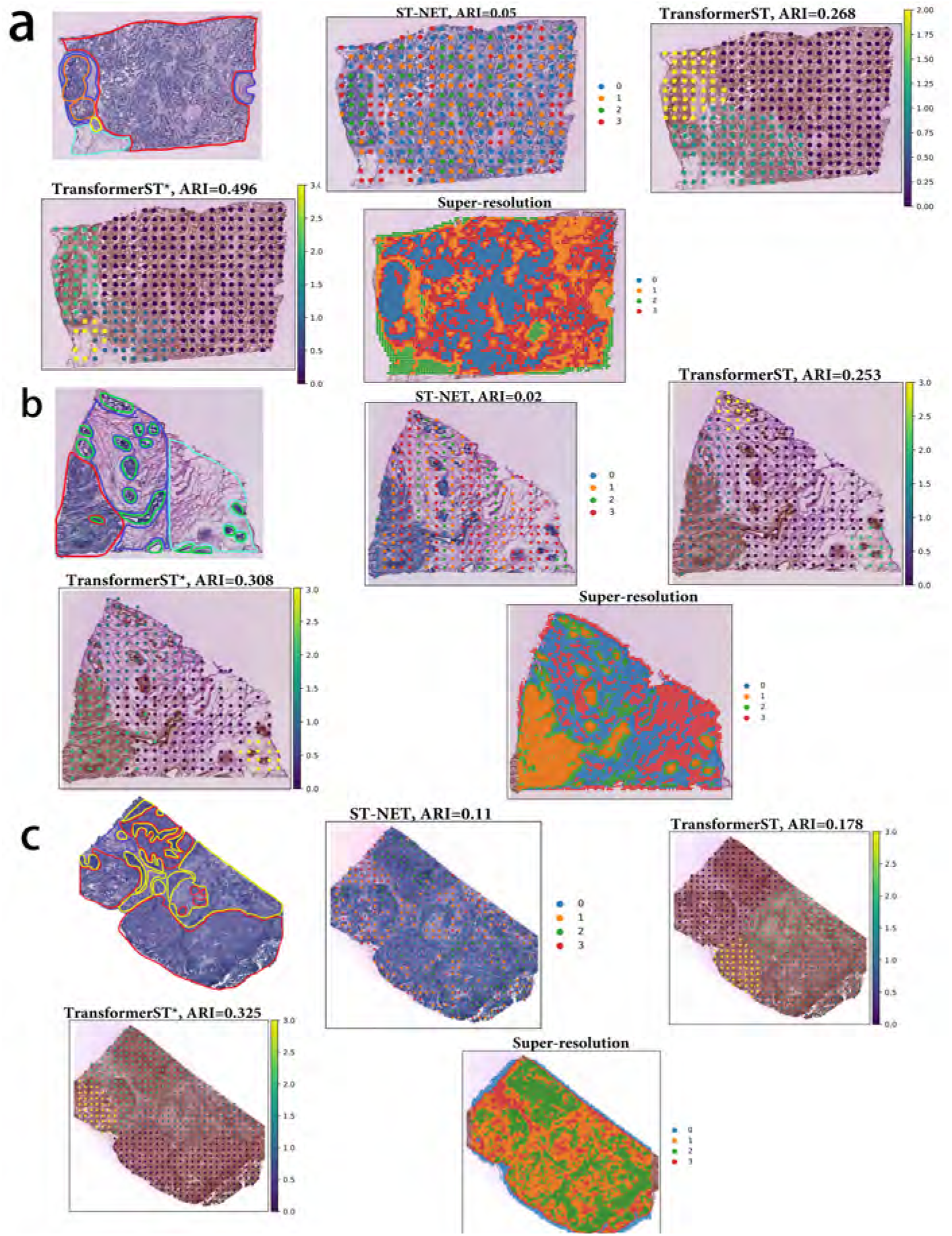
Super-resolved gene expression prediction with breast cancer data. a, Tissue type assignments and nearly single cell super-resolution using A1 section. b, Tissue type assignments and nearly single cell super-resolution using B1 section. c, Tissue type assignments and nearly single cell super-resolution using E1 section.

**Fig. 6.**
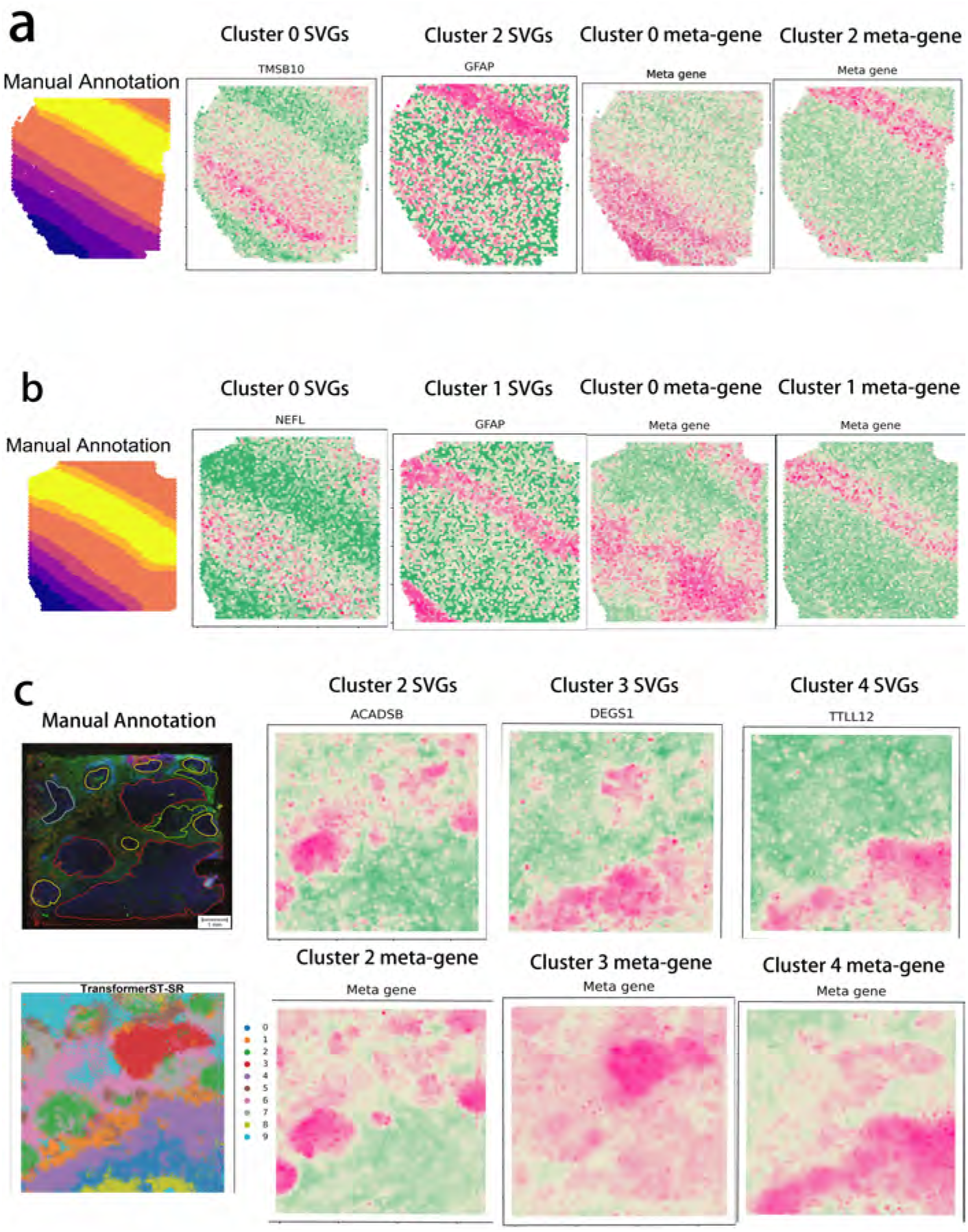
Spatial variable genes (SVGs) and meta gene detection.a, SVGs and corresponding meta genes for cluster 0 (TMSB10, TMSB10+MBP-MT-CO2), cluster 2 (GFAP, GFAP+SNORC-TMSB10+CDT3-MBP) in brain tissue slice 151508 at spot resolution. b, SVGs and corresponding meta gene for cluster 0 (NEFL, NEFL+SCGB2A2-HPCAL1), cluster 1 (GFAP, GFAP+MT1G-FTH1+AQP4-CALM2+CST3-MBP) in brain tissue slice 151509 at spot resolution. c, SVGs and corresponding meta gene for cluster 2 (ACADSB, ACADSB+NME2-MUC1+ATP5MPL-CD74+LAPTM4B-CRIP1), cluster 3 (DEGS1, DEGS1+RPS18-CXCL14+AGR2-MGP+CSTA-NEAT1), and cluster 4 (TTLL12, TTLL12+HMGN2-MALAT1+KRT8-SLC9A3R1) in IDC sample at nearly single cell resolution. Pathologist annotated different regions in different colors (IC outlined in red, carcinoma in yellow, benign hyperplasia in green, unclassified tumor in grey).

### 2.1 Tissue Type Identification at Original Resolution

#### Tissue identification in human dorsolateral prefrontal cortex Visium data

Recently, the LIBD human dorsolateral prefrontal cortex (DLPFC) data were acquired with 10x Visium platform. The whole dataset sequenced 12 tissue samples with manual annotations of six cortical layers and white matter for each sample. The manual annotations are provided by the original study [37] and allows to evaluate the performance of tissue type identification at spot resolution. We evaluated the tissue type identification of TransformerST compared with stLearn, Mclust, Kmeans, Louvain, Giotto, BayesSpace, CCST, STAGATE and SpaGCN. We used the adjusted Rand index (ARI) to quantify the similarity between ground truth and clustering results[37].

The clustering accuracy (ARI) of sections 151672 and 151508 are shown in Fig. 2a and Fig. 2c. Comparing with the baseline methods, TransformerST could learn the dynamic graph representation between spatial gene expression and spatial neighbors. Specifically, the proposed method were implemented using the top 3000 HVGs, other comparison methods such as BayesSpace and SpaGCN used 15 PCs from top 3000 HVGs. Gitto, CCST, STAGATE and StLearn used the recommended parameters in the previous papers. The proposed method could take advantage of the highly expressive gene and spatial dependence of neighboring embedding to achieve the highest tissue identification performance of both samples. Fig. 2a shows, for section 151672, TransformerST, Gitto, STAGATE and SpaGCN revealed spatial gene expression patterns better accord with manual annotations, the ARI is 0.687 for TransformerST, 0.573 for Gitto, 0.561 for STAGATE and 0.565 for SpaGCN. The visual difference among these results are not significant. BayesSpace, Mclust and CCST also provided decent results (ARI is 0.439 for BayesSpace, 0.479 for Mclust and 0.427 for CCST) and outperformed Louvain, StLearn and Kmeans. In Fig. 2c, for section 151508, TransformerST had the highest clustering accuracy and provided distinct layers of clusters (ARI is 0.592). CCST and STAGATE outperformed other methods but provided a worse performance than TransformerST.

The remaining clustering results with all 12 DLPFC samples are shown in Fig. 2b. TransformerST achieved the best performance with mean ARI (0.564). Compared with the second performer STAGATE with mean ARI (0.502), TransformerST increased the tissue identification performance by 12.4%. The difference between BayesSpace, ccST and SpaGCN is not significant. Additionally, the runtime of TransformerST at spot resolution are comparable to other clustering methods for spot level annotation, which uses 6.5 mins with 3000 HVGs and 3 mins for 200PCs. (Table 1). Detailed experimental results with all 12 DLPFC samples are shown in supplementary material.

**Table 1.** Computational time for tissue type identification with LIBD human dorsolateral prefrontal cortex

| Method | Runtime/mins | GPU/CPU |
| --- | --- | --- |
| TransformerST-3000 HVGs | 6.5 | GPU |
| TransformerST-200 PCA | 3 | GPU |
| BayesSpace | 21 | CPU |
| stLearn | 0.5 | GPU |
| SpaGCN | 2 | GPU |
| CCST | 3 | GPU |
| STAGATE | 7 | GPU |
| Gitto | 17 | CPU |

These results further demonstrate the superiority of TransformerST to explore the spatial expression patterns and provide clear cluster difference between brain layers.

#### Tissue identification in mouse lung Visium data at spot resolution

To further assess the performance of TransformerST in tissue identification, we performed Visium experiments on four slices of mouse lungs. Single-cell suspension processed side-by-side was subjusted to scRNA-seq experiment and utilized to deconvolute the Visium data.

Pathologist then identified regions of interest (airways and blood vessels) according to the histology images. Airways were defined in line with the deconvoluted proportion of club cells within each slice. Pathologist manually set the thresholds in each slice to match the selected spots with the histological airways. Spots were marked as airways if the proportion of club cells were above the threshold (top 20% for slice A1, top 20% for slice A2, top 10% for slice A3, and top 10% for slice A4). Blood vessels were defined in consist with the blood vessels regions in the histology image. A random trees pixel classifier using QuPath (version 0.2.3) with downsample = 16 was trained to estimate the probability of blood vessels within each spot of all slice samples. All the training samples of the random trees pixel classifier came from the manual annotation of slice A1. Then, pathologist used the threshold 0.5 to select the blood vessels (Fig. 3b and Fig. 3c).

After defining these histological structures, TransformerST was utilized to reveal the internal heterogeneity within visually homogeneous blood vessel and airway tissue regions. The cluster numbers of all comparison methods were set to 4. Fig. 3a shows, for the first slice sample, SpaGCN, STAGATE and StLearn were able to distinguish the airways, but failed to identify the tissue region of blood vessels. Surprisingly, BayesSpace failed to identify the significant tissue types such as blood vessel and airway (Fig. 3a). Other comparison methods such as Mclust, Kmeans, CCST, Louvain had worse performance, which are contrary to the manual annotation (Fig. 3b). Gitto could identify the major tissue types, but its result are very noisy. The most interesting finding is that TransformerST is able to identify the whole blood vessel regions and provide a more robust signal with detailed textural features (Fig. 3a).

Moreover, we used the club cell tissues to evaluate the performance of TransformerST. As shown in Fig. 3a, for the first slice sample, TransfromerST, SpaGCN, Gitto, STAGATE and StLearn were able to identify the club cell regions, an indicator of airways. We observed that the spatial expression patterns of club cells between the clusters were largely in line with the clinical annotations (Fig. 3b). BayesSpace, CCST and non-spatial methods (Mclust, Kmeans and Louvain) failed to detect the spatial patterns of club cell structures. Comparing these results, it could be seen that spatial expression patterns acquired by TransformerST better reflect the the club cell structures with detailed information in the boundaries.

The relative performance remains the same for the second slice sample (Fig. 3c), TransformerST, StLearn, Gitto, STAGATE and SpaGCN were able to identify the heterogeneity within club cells structure (Fig. 3d). However, as shown in Fig. 3d, these methods aside TransformerST revealed stantial noise and lack of clear spatial difference between club cells. BayesSpace, Mclust, Louvain, CCST and Kmeans provided worse performance which violates the biological interpretation. The existing methods are not applicable for mouse lung tissue identification. TransformerST could identify the spatial patterns with histology image and provided finear details of manual annotations (Fig. 3c).

### 2.2 Spatial Transcriptomics Super-Resolution at Enhanced Resolution

#### Tissue identification and super-resolution in melanoma ST data at subspot resolution

We evaluated the subspot super-resolution performance with the publicly available melanoma ST data which was annotated and described in Thrane et al [14]. The manual annotation of melanoma, stroma and lymphoid regions (Fig. 4a) were included to evaluate the performance of the TransformerST. Similar to manual annotations, we set the cluster number to 4. As the histology image is missing, both BayesSpace and TransformerST could enhance the resolution of ST expression to subspot resolution. We show the tissue identification results of the proposed method in both spot and subspot resolution in Fig. 4a and Fig. 4b. Comparison of the results of TransformerST with those of other methods (Mclust, Kmeans, Louvain, Gitto, SpaGCN, CCST, STAGATE and BayesSpace) confirms that TransformerST reveals similar patterns to the manual annotation.

Specifically, the melanoma tissue could be divided into two types, central tumor region and outer of the mixture of tumor and lymphoid tissue. Surprisingly, only TransformerST was able to identify the lymphoid regions at original resolution (Fig. 4a). The results of comparison methods could not identify lymphoid regions at the original resolution. The tissue identification results at enhanced resolution are in line with the finding that TransformerST identifies lymphoid region in the tumor border with a higher resolution (Fig. 4b). In accord with recent study, BayesSpace and STdeconvolve also identified the lymphoid regions to the tumor at the enhanced resolution (Fig. 4b). The results of this study indicate that all the comparison methods could identify the heterogeneity between border and center of tumor but fail to identify lymphoid tissue at original resolution. TransformerST, STdeconvolve and BayesSpace provided enhanced resolution of tissue structures which makes it possible to identify the lymphoid tissue. The observational results suggest TransformerST provides higher resolution and robust tissue identification results at both original and enhanced resolution. Detailed experimental results with enhanced resolution of three methods are shown in supplementary material.

#### Tissue identification and super-resolution in IDC Visium data at nearly single cell resolution

We evaluated the nearly single cell super-resolution performance with the IDC Visium data with immunofluorescence staining for 4,6-diamidino-2-phenylindole (DAPI) and T cells staining CD3 in [28]. Pathologist identified regions of predominantly invasive carcinoma (IC), carcinoma in situ and benigh hyperplasis were included to evaluated the clustering accuracy at spot resolution (Fig. 4d). Similar to the manual annotations, we clustered the IDC sample into 5 clusters at spot resolution. We used ARI to evaluate the clustering accuracy at spot resolution. The results of the clustering experiment at original resolution indicate that TransformerST achieves the best clustering accuracy with ARI of 0.493 (Fig. 4c). The ARI is 0.423 for BayesSpace against 0.369 for SpaGCN, 0.357 for Mclust and 0.274 for Gitto. However, some comparison methods did not improve the clustering performace (ARI is only 0.257 for StLearn, 0.234 for STAGATE, 0.208 for CCST, 0.151 for Louvain and 0.101 for Kmeans).

We further enhanced the spatial transcriptomics resolution to show the biological relevance with TransfromerST, STdeconvolve and BayesSpace (Fig. 4e and 4f). In accord with the BayesSpace paper [28], we set the cluster number *k* = 10. As shown in Fig. 4e and 4f, TransformerST could identify four clusters (0,3,4,8) related to predominantly IC, one cluster (2) related to carcinoma regions, one cluster (7) identify the benign hyperplasia regions. And clusters (1,5,6,9) are related to the unclassified regions. The result of ByesSpace was in consist with previous report in [28]. However it is hard to quantitatively evaluate the cluster accuracy at enhanced resolution. The results of three methods show the spatial heterogeneity among tumor which is inaccessible to histopathological analysis. However, We saw the visual difference between carcinoma and benigh hyperplasia regions via TransformerST compared to BayesSpace and STdeconvolve. TransformerST exhibited the spatial organization more similar to manual annotations. BayesSpace could only increase the IDC data to subspot resolution, TransformerST could predict the heterogeneity within each tissue at nearly single cell resolution. STdeconvolve revealed the proportion of each cell type, but failed to identify the location of cell patterns within each spot. The runtime of TransformerST at enhanced resolution are comparable to other methods for gene expression reconstruction, which uses 29 mins. (Table 2). TransformerST provides a more efficient approach to identify the super-resolved tissue microenvironment than BayesSpace and STdeconvolve. Detailed experimental results with enhanced resolution of IDC samples are shown in supplementary material.

**Table 2.** Computational time for super-resolved gene expression reconstruction with IDC sample

| Method | Runtime/mins | GPU/CPU |
| --- | --- | --- |
| TransformerST-3000 HVGs | 29 | GPU |
| BayesSpace | 200 | CPU |
| STdeconvolve | 54 | CPU |

### 2.3 Enhanced Gene Expression Prediction at nearly Single-Cell Resolution

#### Enhanced Gene expression prediction at nearly single cell resolution in breast cancer data HER2+

To predict gene expression at nearly single cell resolution using histology image, we used two experiments to evaluate the tissue identification and tissue super-resolution performance, the leave-one-out evaluation (36 fold) and single section evaluation.. We used the HER2+ breast cancer data which includes 36 tissue sections from 8 patients to demonstrate the performance of gene expression prediction and super-resolution. Specifically, for leave-one-out evaluation, 32 sections were included to train the tissue identification and super-resolution model and the remaining section were used for evaluation. The results of leave-one-out are represented as TransformerST. We also showed the clustering results of the TransformerST using single tissue section, which is marked as *TransformerST*\*. We further evaluated the superresolution performance at nearly single cell resolution, which is represented as Super-resolution.

Manually annotation of 3 tissue sections were included for evaluation of clustering accuracy. We compared the proposed method with ST-NET [38] for gene expression prediction using three tissue sections in Fig. 5. The ST-NET ignores the spatial relationship between spots and showed worse gene prediction performance. Both the leave-one-out evaluation and single section evaluation yielded higher correlation with biological interpretation. From Fig. 5 we can see that TransformerST increased the clustering accuracy (ARI) for three sections (A1, B1, E1), which is much higher than those predicted by ST-NET. For example, as shown in Fig. 5a, for sample A1, TransformerST outperformed ST-NET with higher clustering accuracy (ARI is 0.268 for TransformerST, but 0.05 for ST-NET). *TransformerST\** could predict much higher ARI (ARI=0.496) than TransformerST and ST-NET. This result might be explained by the fact that there are strong gene expression difference among patients, the single section evaluation (*TransformerST*\*) achieves better tissue identification performance (Fig. 5). The relative performance remains the same for sample B1 (Fig. 5b). *TransformerST*\* outperformed ST-NET and TransformerST with the highest clustering accuracy (ARI is 0.308 for TransformerST, 0.253 for TransformerST and 0.02 for ST-NET). Fig. 5c shows superiority of *TransformerST\** (ARI=0.323) over TransformerST (ARI=0.178) and ST-NET (ARI=0.11).

The enhanced single cell resolution results further demonstrate TransformerST could predict the biological meaningful patterns as in the manual annotations. While it is bard to estimate the ARI for the super-resolution result, the study is visually consist with the manual annotations by pathologists in the spatial domain (Fig. 5).

### 2.4 Meta Gene and SVGs Analysis with DLPFC and IDC Samples

To further demonstrate that TransformerST could explore the biological relevance, we detected the spatial variable genes and meta genes for LIBD human dorsolateral prefrontal cortex (DLPFC) data and IDC sample. As shown in Fig. 6a and Fig. 6b, SVGs and their corresponding meta gene show similar spatial patterns for human DLPFC samples at spot resolution. For example, TMSB10 is enriched in cluster 0 of tissue sample 151508. The combination of meta gene (TMSB10+MBP-MT-CO2) shows the strengthened spatial patterns in the neighboring regions. GFAP is enriched in cluster 2 of tissue sample 151508, its corresponding meta gene is GFAP+SNORC-TMSB10+CDT3-MBP, which is also spatilly correlated with the SVGs of cluster 2 in the histology image.

TransformerST also detected a single SVG to mark the corresponding spatial domain in tissue sample 151509. NEFL is enriched in cluster 0 with the visually corresponding meta gene defined as NEFL+SCGB2A2-HPCAL1. GFAP is enriched in cluster 1 which is visually consist with its meta gene (GFAP+MT1G-FTH1+AQP4-CALM2+CST3-MBP). Both SVGs and meta gene show similar spatial patterns in the histology image (Fig. 6b). The experimental results with different tissue samples and different cluster domains demonstrate TransformerST could mark specific gene expressed regions for different cluster domains.

To illustrate how TransformerST works for different tissue samples, We detected the spatial variable genes and meta gene for IDC sample at nearly single cell resolution. As shown in Fig. 6c, TransformerST detected single SVGs (ACADSB) for cluster 2. Its corresponding meta gene was defined as ACADSB+NME2-MUC1+ATP5MPL-CD74+LAPTM4B-CRIP1. TransformerST detected DEGS1 SVG for cluster 3, which accords with its meta gene DEGS1+RPS18-CXCL14+AGR2-MGP+CSTA-NEAT1 visually. TTLL12 is enriched in cluster 4 with its corresponding meta gene as TTLL12+HMGN2-MALAT1+KRT8-SLC9A3R1.

The detection results of meta gene and SVGs reflect that TransformerST is able to identify the heterogeneity among spatial domains and predict the boundaries not annotated by pathologists. These results demonstrate TransformerST could better explore the spatial patterns with graph transformer network.

## 3 Discussions

In the study, we propose a novel Transformer based method for integrating the gene expression, the spatial location and histology image (if available). The proposed method called TransformerST is the first time to enhance the spatial transcriptomics to nearly single cell resolution without single cell RNA-seq reference. Different from most of the existing spatial transcriptomics analysis methods, TransformerST does not require the linear PCA preprocessing and guarantees the nonlinear learning of spatially distributed tissue structure of multimodal data (i.e. ST and 10x Visium). The adaptive graph transformer model with multi-head attention makes it possible to associate multimodal graph representation and reveal how the heterogeneity map shapes the tissue function dynamics. With the help of a cross-scale internal graph network, TransformerST enables the effective and efficient analysis of super-resolved tissue microenvironment at nearly single cell resolution. We evaluate the performance of TransformerST with several datasets generated using diverse spatial transcriptomics technologies. Compared with the state-of-the-art methods, TransformerST is able to identify the tissue clusters at both spot and nearly single cell resolution. TransformerST overcomes the limitation of low resolution of current spatial transcriptomics technology and provides an efficient way to explore the spatial neighboring relationship. The experimental results demonstrate the importance of regional heterogeneity and the corresponding intrinsic structural-function relationship within tissue dynamical microenvironment. TransformerST could lower the computation complexity and memory usage than existing methods.

Although the tissue type identification research is an important topic in current spatial transcriptomics analysis, from the experimental results, we could see that most of the state-of-the-art methods fail to estimate the cell heterogeneity within each cell type. We expect TransformerST could help to provide better resolution of spatial transcriptomics data analysis. TransformerST could achieve super-resolved resolution of single cell per subspot without the requirement of additional single cell RNA-seq reference. However, TransformerST could also be easily adapted to incorporate additional single cell reference for deconvolution task. With the downstream analysis such as SVGs and meta gene analysis, TransformerST shows the similar biological tissue patterns to manual annotations.

While TransformerST focuses on the ST and Visium platform, it could be easily applied to other platforms with slight modification. In summary, TransformerST provides an effective and efficient pipeline for various unsupervised spatial transcriptomics analysis such as tissue identification, super-resolved gene expression reconstruction and gene prediction from histology image. For future work, we anticipate to increase the tissue type identification accuracy by estimating the contribution of cell-specific gene expression. We also intend to improve the graph transformer model to explore the heterogeneity of tissue type in different micro-environments.

## 4 Methods

### Data description

TransformerST is evaluated using several public available datasets and one in-house dataset, most of which were obtained via Visium platform. Specifically, the DLPFC dataset includes 12 sections. The number of spots within each section ranges from 3498 to 4789. The area of DLPFC layers and white matter (WM) were manually annotated by pathologists. To reconstruct gene expression at enhanced resolution, we use the publicly available melanoma ST data which was annotated and described in Thrane et al [14]. We show the performance of super-resolution at nearly single cell resolution using IDC Visium data with immunofluorescence staining for 4,6-diamidino-2-phenylindole (DAPI) and T cells staining CD3 in [28]. To predict gene expression at nearly single cell resolution using histology image, we used the HER2+ breast cancer data which includes 36 tissue sections from 8 patients. We also use our in-house mouse lung data to evaluate the performance of TransformerST in tissue identification experiment.

### In-house data preprocessing

For our in-house mouse lung data, 10X Genomics Visium platform were used to perform the ST experiment. After harvesting the mouse lungs, the left lobes were inflated with 1mL of mixture of 50% sterile PBS/ 50% Tissue-Tek OCT compound (SAKURA FINETEK) before being frozen in alcohol bath on dry ice. Until they were processed further, OCT blocks were kept at −80°C Following the 10x Genomics Visium fresh frozen tissue processing protocol, OCT blocks were sectioned at 10*μ*m in thickness and 6.5mm X 6.5mm in size, affixed to the Visium slides, and then stained with hematoxylin and eosin. A fluorescence and tile scanning microscope (Olympus Fluoview 1000) was used to take H&E images, after which the slides underwent tissue removal and library generation per 10x Genomics demonstrated protocol.

Using Space Ranger software (version 1.2.2) from 10x Genomics, each sequenced spatial transcriptomics library was processed and aligned to the mm10 mouse reference genome, with UMI counts summarized for each spot. Tissue overlying spots were identified based on the images in order to distinguish them from the background. When the filtered UMI count matrices were generated, only the barcodes associated with these tissue overlaying spots were kept. Additionally, we manually excluded spots that were not covered by tissue but were yet detected by Space Ranger and further filter the UMI count matrices (slice A1: 32,285 genes x 3,689 spots; slice A2: 32,285 genes x 2,840 spots; slice A3: 32,285 genes x 3,950 spots; slice A2: 32,285 genes x 3,765 spots).

### Public data preprocessing

All Visium samples were generated from 10x Genomics procured from BioIVT:ASTERAND. The remaining melanoma and breast cancer samples were obtained using the ST platform. We use the second replicate from biopsy 1 to detect the lymphoid sub-environment. For all datasets, raw genes expression counts expressed in fewer three spots were filtered and eliminated. Seurat were then introduced to find the top 3000 most HVGs for each spot. The gene expression values are transformed into a natural log scale. We use both histology image (when available) and spatial gene expression to exploit tissue sub-environment at the super-resolved resolution.

### Graph reconstruction for spatial gene expression

TransformerST reconstructs the cell-cell relationship using an undirected graph G(V,E). Each vertex *V* represents the spot and the edge *E* measures the weighted relationships between two vertices. We map each spot back to the histology image and define the corresponding pixel using similar smooth and rescale steps in SpaGCN [30]. The Euclidean distance between vertices is calculated based on the image coordinates. We select top 20 neighbors of each spot to represent the adjacency matrix *A.*

### Transformer based variational auto-encoder representation learning

The transformer based variational auto-encoder is introduced to explore the feature learning capability at both spot and nearly single cell resolution. The neighboring relationship of spatial transcriptomics requires the proposed method to understand the global composition of histology image and corresponding gene expression, which enables it to reconstruct the locally realistic and globally consistent patterns of gene expression. Thus, we use a codebook to represent the perceptually rich gene expression patterns. Together with graph transformer model, the Variational-Transformer architecture could reconstruct the realistic and consistent enhanced resolution spatial gene expression in a conditional setting. The proposed VAE-Transformer model consists of three parts, codebook representation learning, transformer-based reconstruction and conditional synthesis.

#### Learning an effective codebook of gene expression constituents

The aim of the codebook learning is to exploit the the constituents of the spatial gene expression in the form of a sequence. The spatial gene input for the Transformer-based variational autoencoder is stored in *N* × *M* matrix which consists of *N* spots and *M* genes. In addition, we represent the histology image as *H* × *W* with 2 dimensional coordinates. Together with the spatial coordinates, the spatial gene expression could be represented using a spatial collection of codebook entries *z_q_* ∈ *R^*h*×*w*×n_z_^*, where *n_z_* is the dimensionality of codes. Similar to neural discrete representation learning, we propose a convolutional model which consists of an encoder *F* and a decoder *D* to exploit the discrete codebook *Z* = *z_k_*, (*k* = 1,…, *K*). Each item of the coderbook *z_q_* is obtained via the encoder *z* = *F*(*x*) and an element-wise quantization *Q*(.) of each spatial code *z_i,j_*.

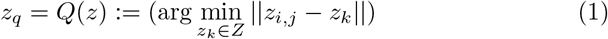

The reconstructed spatial gene expression is defined as

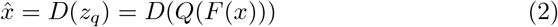

We simply copy the gradient from the decoder to the encoder and train the codebook learning model via an end-to-end loss function

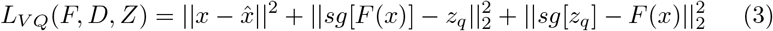

where 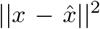 represents the reconstruction loss. *sg*[.] denotes the stop-gradient operation.

#### Learning the spatial gene expression with a conditional Transformer

Instead of straightly representing the quantized encoding *z_q_* = *Q*(*F*(*x*)), we use the conditional transformer model to represent the codebook indices *s*.

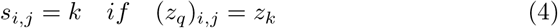

With the learned sequence of learned codebook indices *s*, we could map *s* back to *z_q_* = (*z_s_i,j__*)and reconstruct the original spatial gene expression 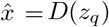.

Then the transformer is used to predict the distribution of next indices in a conditional setting,

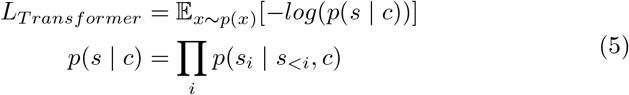

where the condition *c* is defined as the clustering result of graph-transformer model. Finally, the spatial gene expression is reconstructed in a sliding-window manner. In order to accelerate the training process, we crop the spatial gene expression into patches and restrict the length of *s*.

### Adaptive graph-transformer for spatial embedding

The proposed method utilizes the adaptive graph transformer model to embed the spatial relationship of neighboring spots. The proposed method concatenates the gene expression embedding *F*(*x*) and edge weights to cluster each spot. For the downstream analysis, the Graph Transformer layer together with the multi-head attention model is utilized to stack the entire node features. The inputs for the multi-head attention consists of query, key and value. We define the multi-head attention for each edge with each layer *l* as follows,

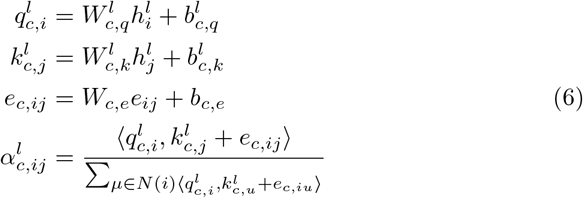

where 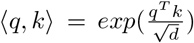 represents the exponential scale dot-product function. *d* is the hidden size of each head. We use the learnable parameters 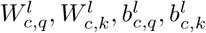 to transform each source feature 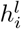 and distant feature 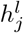 into query vector 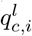 and key vector 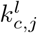. The additional edge feature *e_ij_* is also added into the key vector 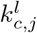.

The message aggregation from *j* to *i* is defined as follows,

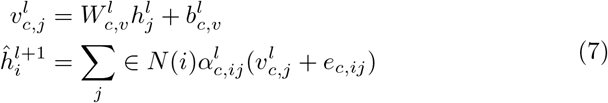

A gated residual connection between layers is adopted to prevent over-smoothing.

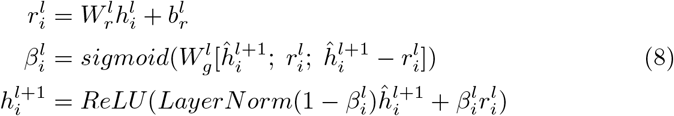

The output of last layer is the averaging of multi-head output

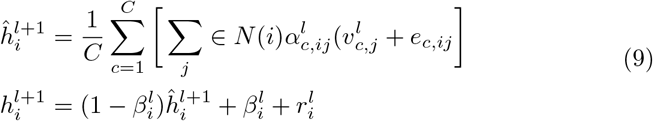

#### Adaptive Graph transformer representation learning

The previous spatial transcriptomics clustering method only considers the spatial information to construct the graph representation. We introduce an adaptive Graph Transformer model to learn the spatial and feature representation of the entire graph, which is defined as follows:

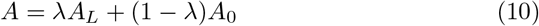

where *A*_0_ is the initial adjacency matrix and *A_L_* is the learned adjacency matrix within each iteration. The initial adjacency matrix is constructed using the *k* nearest neighborhood using the histology image. The adaptive updating mechanism helps to learn the global and local representation of spatial transcriptomics data. The hyperparameter λ is included to balance the trade-off between spatial and feature graph structure.

#### Identifying tissue types with iterative clustering

Based on the outputs of Graph Transformer encoder, the proposed method iteratively identifies the tissue type in an unsupervised manner. The initiation of the proposed method is based on Louvain’s method. The clustering method is divided into two steps. In the first step, we assign a soft cluster type *γ_i,j_* to each spot embedding *z_i_* as follows:

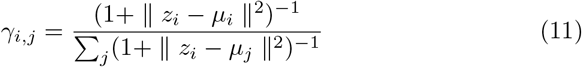

Then we refine the clusters with an auxiliary target distribution *p* based on *γ_i,j_*

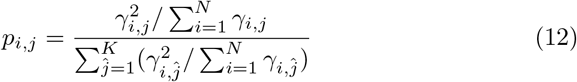

Similar to the previous iterative clustering algorithm in scRNA-seq analysis, the loss function is defined using the KL divergence.

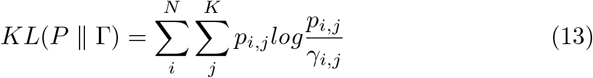

### Reconstructing the super-resolved gene expression at the sub-spot resolution

In order to explore the tissue sub-environment at the enhanced resolution, we segment each spot into nearly single cell resolution with the help of histology image. If the histology is missing in real time applications, we adopt the setting of BayesSpace [28], each spot ST data is segmented into nine subspots and each Visium data is segmented into six subspots. As the ST spots are 100 μm in diameter and Visium are 55 μm in diameter, TransformerST could achieve the super-resolved gene expression at nearly single cell resolution rather than the original mixture of dozens of cells. The proposed super-resolved reconstruction components are divided into twp steps, histology image super-resolution and spatial gene expression reconstruction.

We model the internal cross-scale relationship between each spot and its corresponding neighbors as graph, where each spot patch is the vertex and the edge is the weighted connection of two spots in different resolutions. The proposed method consists of two parts, graph construction and patch aggregation. TransformerST dynamically reconstructs the cross-resolution graph by searching the *k* nearest neighboring spots in the downsampled resolution. With the mapping function, we could obtain the *k* nearest neighboring spots patch at original resolution. Thus, the reconstructed graph provides *k* spot mapping pairs of low and high resolution. After that, we employ the patch aggregation model to aggregate *k* spot patches conditioned on the similarity distance. Due to the limitation of current spatial transcriptomics technology, we could not get the ground truth data at the enhanced resolution. We assume that the spatial gene expression at the spot resolution is the averaged mixture of its corresponding single cell segments. Instead of straightly calculating the reconstruction loss at the enhanced resolution, we average the single cell components into spot to guide the training process.

#### Graph reconstruction

We first downsample the spot gene expression by the factor *η*. *η* is set to nine for ST platform and six for Visium platform. The downsampled spatial gene expression is denoted as *X*_*L*↓*η*_. After that, we find the *k* neighboring patches in graph at low and high resolution. To obtain the *k* neighboring patches, we extract the embedded features by the encoder model of graph-Transformer and variational encoder. For each spot, we search its *k* neighboring spot in *X*_*L*↓*η*_ and *l* × *l* patches in *X_L_* with Euclidean distance. We search the similar spot in *X*_*L*↓*η*_ rather than *X_L_,* we could lower the search space by *η*^2^.

#### Patch aggregation

We weight the *k* neighboring patches on the similarity distance and aggregate the enhanced gene expression as

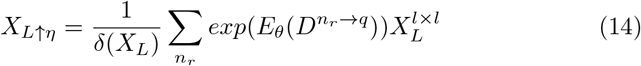

where *δ*(*X_L_*) = ∑*_n_r__ exp*(*E_θ_*(*D*^*n_r_*→*q*^)) denotes the normalization factor. *E_θ_*(*D*^*n_r_→*q**^) is used to estimate the aggregation weight for each neighboring patches.

### Spatially variable meta genes detection

We are interested in the detection of spatially variable meta gene within each tissue type. The spatially variable meta gene expression could be divided into two steps. The first step is to detect the spatial variable gene (SVGs) in the target tissue type but not high expressed in its neighbors. The number of neighbors is set to 10 in the experiments. We select a nontarget tissue type domain using the threshold 50%. Specifically, if more than 50% spots of a nontarget tissue type domain are in the neighboring set, we will define that tissue type as neighboring tissue type domain. Genes with FDR-adjusted *P* value < 0.05 are defined as spatial variable genes.

The second step is to detect the spatially variable meta genes. For each specific tissue type, we are interested to detect the set of multiple genes shows an enriched expression patterns. The expression of meta gene is defined as follows,

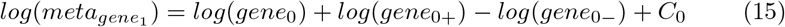

where *C*_0_ is a constant to make meta gene expression non-negative. *gene_0_+* is selected with the higher expressed genes and the smallest FDR-*P* value in the target tissue type. Similarly, *gene*_0+_ is included with the higher expressed genes and the smallest FDR-*P* value in the control tissue type. The output of meta gene detection is obtained iteratively,

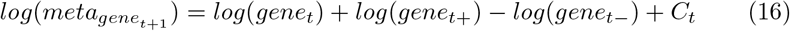

### Moran’s I and Geary’s C statistics for the evaluation of SVGs

Moran’s I metric is a correlation coefficient to measure the overall spatial autocorrelation of a dataset. We define the Moran’s I for the given spatial variable gene as follows,

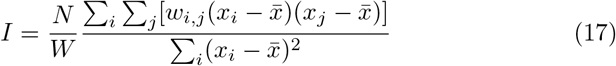

where *x_i_* and *x_j_* are gene expressions of spot *i* and *j*, 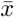 is the mean gene expression. *w_i,j_* is the spatial weight.

In addition, we also use another commonly used statistic model Geary’s C, which is defined as,

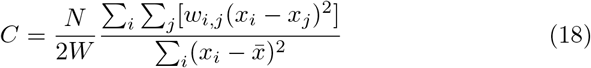

The values of Geary’s C should be similar to Moran’s I for each specific gene expression.

## 5 Data Availability

We use several publicly available data which could be acquired using the following websites or accession numbers: (1) LIBD human dorsolateral prefrontal cortex data (DLPFC) (http://research.libd.org/spatialLIBD/); (2) Melanoma ST data (https://www.spatialresearch.org/wp-content/uploads/2019/03/ST-Melanoma-Datasets_1.zip); (3) Human epidermal growth factor receptor(HER) 2 amplified (HER+) invasive ductal carcinoma (IDC) sample [28]; (4)HER2+ breast cancer data [38]

## Notes

### Competing Interest Statement

The authors have declared no competing interest.

